# Key gut microbiota components and functions in an aquatic keystone species across diets assessed by metaproteomics

**DOI:** 10.1101/2024.11.06.622251

**Authors:** Thibaut Dumas, Olivier Pible, Lucia Grenga, Davide Degli Esposti, Nicolas Delorme, Olivier Geffard, Arnaud Chaumot, Jean Armengaud

## Abstract

The gut microbiota plays a crucial role in maintaining host fitness and modulating contaminant toxicity-related responses. However, information on how the gut microbiota of sentinel species responds to environmental factors is limited. In this study, we characterized the gut microbial community and its functions under normal, contaminant-free conditions by examining the effects of different diets over a 10-day period (alder leaf, carrot, spinach, and protein-rich granules) on the amphipod *Gammarus fossarum*, commonly used in bioassays for ecotoxicity assessment of contaminated rivers. Metaproteomic analysis of intestine samples enabled taxonomic characterization of the gut microbiota from this millimetric animal, assignment of biological functions to each microbial entity, and functional analysis of host proteins. The most abundant microbes detected in the gut belong to 37 bacterial and 5 fungal genera. Functional analyses of host and microbial proteins revealed complementary metabolic activities, allowing the degradation of complex polysaccharides such as cellulose and chitin. Diet was found to shape microbial community structure, with foodborne microorganisms strongly influencing structural changes during short-term feeding in amphipods. These microorganisms remained viable post-ingestion and contributed to food digestion. Functional stability was maintained across different diets, although the protein-rich granules diet induced functional shifts in both the host and its microbiota, reflecting their adaptation to a novel nutrient source. Finally, we identified a core microbiota driving key gut functions, less affected by dietary variations. These findings are significant for future ecotoxicological and biomonitoring investigations, leveraging the microbiomes of these sentinel animals as pivotal targets.

## Introduction

The digestive systems of animals harbor a rich array of microbial communities that typically act as mutualistic symbionts, contributing to host fitness ^1^. Abundant research data extensively document how the gut microbiota assist their hosts in pivotal functions including digestion, nutrient processing, immune function, disease resistance, and metabolism of xenobiotics ^2^. Collectively, the microbiota and the molecular products of their diverse activities, including proteins, metabolites, transcripts, chromosomes and mobile genetic elements, whether contained within cells or dispersed throughout their surrounding environment, are commonly referred to as the “microbiome” ^3^.

While shotgun metagenomics enables to assess the taxonomic composition and gene repertoire of microbial communities ^4^, functional-focussed omics approaches such as metatranscriptomics, metaproteomics ^5^ and metabolomics ^6^ can be employed to gain deeper insights into microbiome functions. Metaproteomics relies on the extraction of proteins, their subsequent proteolysis into peptides by trypsin, and their analysis by liquid chromatography coupled to tandem mass spectrometry (LC-MS/MS) ^7^. MS/MS spectra are assigned to peptides using protein sequence databases sourced from either metagenomics data, reference genomes or transcriptomes. Metaproteomic workflows are capable to assign a functional and taxonomic annotation to each peptides or identified proteins, while also quantifying the relative abundance of each taxon and its associated functions ^8^. Furthermore, metaproteomics allows to gain information on the proteins produced and released by the host ^9,10^, thereby shedding light on the intricate interactions between microorganisms and their host. A workflow has been described wherein taxonomic units are established through proteotyping, followed by the confident identification of proteins and their functions using a sample-specific database, thus providing species resolved granularity ^11^.

Previous studies have established that the gut microbiota of animals can be sensitive to a wide range of factors. While most research has focused on vertebrates, particularly mammals ^12^, a few studies have demonstrated the sensitivity of the gut microbiota from aquatic invertebrates to external or intrinsic factors such as diet ^13^, environmental conditions ^14,15^, habitat ^16^, pathogens ^17,18^, chemical exposures ^19^, or sex ^20,21^ and developmental stage ^22,23^. Although the selection of aquatic invertebrates for study is often driven by economic rationale (e.g. shrimp aquaculture), there is a critical need to investigate the gut microbiota of ecologically relevant species by integrating such factors to better understand the role of their microbiota and host-microbiota interactions.

Gammarids constitute a diverse group of amphipod crustaceans, comprising over 200 species distributed across inland and coastal waters of the Holarctic ^24^. They rank among the most abundant macroinvertebrates in running rivers in terms of biomass, thus gammarids represent keystone species in aquatic ecosystems ^25,26^. Acting as shredders and detritus feeders, they actively contribute in organic matter cycling (e.g., litter breakdown processes), and they serve in return as prey for secondary consumers ^27^. Beyond their ecological significance, European *Gammarus* species, such as *G. fossarum* and *G. pulex*, are of ecotoxicological importance due to their sensitivity to a wide range of chemical stressors ^28^. Protocols using the sentinel species *G. fossarum* have been devised for toxicity testing under laboratory conditions ^29^ and field assessment via active biomonitoring using caging ^30^. Improving our knowledge of the gut microbiota dynamics of *Gammarus* species holds relevance for both ecological and ecotoxicological purposes. Gouveia et al. developed in a proof of concept study a specific workflow combining proteogenomics and metaproteomics to analyze the microbiome of this millimetric animal of environmental interest but for which the genome is not yet sequenced ^31^.

This study aims to characterize the taxonomy and the global functionality of the gut microbiota of *G. fossarum*, as well as its change under different feeding diets. Although several studies have shown that *G. fossarum* exhibits a preference for foods colonized by fungi and bacteria which enhance food palatability ^32–34^, none has investigated the impact of diet on the shaping of its gut microbiota. Here, four distinct diets were tested, including alder leaves, serving as the reference diet representative of river litter, along with carrots, spinaches and protein-enriched granules intended for shrimp farming. The choice of these foods lies in the difference in their nutritional content mainly in terms of polysaccharides (e.g. pectin, cellulose, hemicellulose and lignin), free sugars (e.g. sucrose, glucose, xylose and fructose), proteins (animal and vegetal) and lipids (e.g. phospholipids). Our metaproteomics approach leverages a sensitive and robust LC-MS/MS data acquisition method, coupled with a multi-step search strategy and a tailored, sample-specific database. In this study, we demonstrate that this approach is particularly effective for complex samples with minimal biological material and a low-abundance microbial signal comparing to host signal. Metaproteomics has been successfully applied to millimetric gut samples of sentinel species, offering a comprehensive understanding of the host-microbiota functional networks that underpin essential physiological functions and their interactions with the environment. Key findings include the identification of a core microbiota and the significant role of foodborne microorganisms in shaping the gut microbiota of *G. fossarum*.

## Material & Methods

### Animals

Organisms were collected on a former watercress site (45^◦^57′25.8′′N 5^◦^15′43.6′′E) which shelters the reference source population of *Gammarus fossarum* used in the INRAE laboratory since more than ten years. Organisms were harvested by kick sampling using a net and sieved in the field to target the adult size class. Sampled organisms were quickly transported to the laboratory and kept in 30-L aquaria supplied with continuous drilled groundwater under constant aeration for a two-week period of acclimation to lab conditions. Water conductivity was around 300 µs/cm. All further experiments with different food substrates were conducted in the same water. A 16/8 h light/dark photoperiod was maintained. The temperature was kept at 12°C. During acclimation, organisms were fed *ad libitum* with dried naturally senescent alder leaves (*Alnus glutinosa*) conditioned for around three weeks in water before being delivered to gammarids. Freeze-dried worms (*Tubifex tubifex*) were provided as a dietary supplement twice a week.

### Experimental design and diets

Feeding experiments were carried out during 10 days in 1-L aquaria under the same continuous water renewal, with 30 organisms per aquarium (one aquarium for each diet). To exclude any confounding influence of life stage and sex on gut microbiota composition, only adult males with a body size around 11 mm (+/- 0.5 mm) were selected for feeding experiments ^35^. Four diets were tested in *ad libitum* conditions: (i) conditioned alder leaves (similar to acclimation period), (ii) frozen chopped spinach (purchased from an aquarium food supplier), (iii) slices of fresh carrots (bought in an organic farm store) kept frozen in the lab for a couple of weeks before the experiment, and (iv) commercial shrimp food granules (dried pellets from an aquarium food supplier) advertised as rich in marine proteins. At the end of the ten-day exposure period, nine organisms were randomly selected in each aquarium for dissection. Intestines were collected under stereomicroscope, pooled in threes in Eppendorf tubes and frozen directly in liquid nitrogen. A sample of remaining food was also frozen for further analyses of microbial communities present in the food ingested by gammarids. Because it is very difficult to obtain only the gut material from the millimetric amphipods, metaproteomics was performed on individual intestine dissected from each animal (three replicates per diet of three pooled intestines, i.e. nine animals used per diet), thus a large quantity of host signal is expected.

### Sample preparation

#### Protein Extraction

Proteins from the pooled intestines and from the food samples were extracted according to a protocol adapted from Hayoun et al. (2019). Briefly, samples were suspended in 60 µL of LDS buffer containing 26.5 mM Tris HCl, 35.25 mM Tris base, 0.5% LDS, 2.5% Glycerol, and 0.13 mM EDTA, complemented with 5% beta-mercaptoethanol. Samples were sonicated for 10 min in an ultrasonic water bath (VWR ultrasonic cleaner). Samples were transferred into 0.5 mL Screw Cap microtubes (Sarstedt, Nümbrecht, Germany) containing 50 mg of beads. Bead beating was performed with a Precellys Evolution instrument (Bertin Technologies, Montigny-le-Bretonneux, France) at 10,000 rpm for 10 cycles of 30 s, with 30 s of pause between each cycle. Samples were centrifuged for 5 min at 16,000× *g* and the resulting supernatants were transferred to new microcentrifuge tubes before incubation at 99 °C for 5 min in a thermomixer (Eppendorf, Hamburg, Germany).

#### Proteolysis

30 μL of protein extract were subjected to SDS-PAGE on a 10-well NuPAGE 4–12% gradient (Invitrogen) for 6 min at 200 V with MES buffer. Gels were stained with Coomassie Blue Safe stain (Invitrogen) and destained overnight with water. The whole protein content from each well was fractionated in two polyacrylamide bands per sample and processed for further destaining, iodoacetamide treatment, as described previously ^37^. Proteins were subjected to proteolysis with trypsin (Roche) using 0.01% ProteaseMAX surfactant (Promega). The resulting peptides were acidified with trifluoroacetic acid (final concentration 0.5%) and were transferred in a glass vial and stored at 4°C prior to analysis. Peptide concentrations were determined using the Pierce Quantitative Peptide Assays & Standards kit (Ref. 23290; Thermo Fisher Scientific) according to the manufacturer’s instructions.

### Liquid Chromatography - Tandem Mass Spectrometry acquisition

The peptide samples were analyzed with an Orbitrap Exploris 480 (Thermo Scientific) tandem mass spectrometer coupled to a Vanquish Neo UHPLC (Thermo Scientific). Peptides were desalted on a reverse-phase PepMap 100 C18 μ-precolumn (5 mm, 100 Å, 300 mm i.d. × 5 mm, Thermo Scientific) and separated on a 50 cm EasySpray column (75 mm, C18 1.9 mm, 100 Å, Thermo Scientific) at a flow rate of 0.250 μL/min using a 90 min gradient (5%–25% B from 0 to 85 min, and 25%–40% B from 85 to 90 min) of mobile phase A (0.1% HCOOH/100% H_2_O) and phase B (0.1% HCOOH/100% CH_3_CN). Data were recorded in data-dependent acquisition mode with a full mass scan from 375 to 1500 (m/z), a MS resolution of 120,000 and a MS/MS resolution of 15,000. The top 50 precursor ions detected at each scan cycle, with potential charge state 2^+^ or 3^+^, were sequentially selected and subjected to fragmentation. MS/MS scanning was initiated with an intensity threshold of 10,000, an isolation window set at 1.1 Da, and a dynamic exclusion time of 20 s.

### Metaproteomics data processing

The Mascot Daemon 2.6.1 search engine (Matrix Science) was utilized to interpret MS/MS spectra into peptide sequences through a multi-round search procedure derived from previously described protocols ^31,38^. The specified search parameters for the first round included full trypsin specificity, a maximum of one missed cleavage, mass tolerances of 3 ppm on the parent ion and 0.02 Da on the MS/MS. Additionally, carbamidomethylated cysteine (+57.0215) was set as a fixed modification, while oxidized methionine (+15.9949) was considered as a variable modification. The queried database was NCBInrS, an in-house assembled subset of the NCBInr database with one representative annotated genome per species belonging to the superkingdoms Bacteria, Archaea and Eukaryota as initially described ^9^. This database includes 94,176,939 protein sequences from 50,995 taxa, totalling 39,636,215,241 amino acid residues. For each sample, the Taxon-Spectrum Matches (TSMs) as first defined in Pible et al. (2020) to quantify taxa with their assigned MS/MS spectra and the number of taxon-specific peptide sequences to validate identified taxa were measured for each identified taxon at each possible taxonomical rank. Families were validated based on the number of family-specific peptides and TSMs. The annotated genomes from the NCBInr database, available for species within the identified families, were compiled into a new database and queried in a second search round. The same settings were used, except for the maximum missed cleavages and the mass tolerance on the parent ion set to 2 and 5 ppm, respectively. Genera were identified, and the genomes of all their corresponding species available in NCBInr were compiled into the final database. The final database was completed with an RNA-seq informed protein database specific for the host ^40^. The protein sequences identified in the food samples after the second search were added to the final database to ensure it was as representative as possible of the proteins present in the gut sample. The third search was performed with the same settings to identify peptides and proteins with a false discovery rate (FDR) of 1%. The FDR was estimated with a search against the corresponding decoy database with the Mascot engine. Occurrence of a genus in a diet condition was validated if the number of TSMs associated to genus-specific peptides was superior to 5 in at least two replicates. The abundance of each taxa at the genus level was expressed in specific TSMs (speTSMs) corresponding to spectral counts of peptides only reported for a given genus. Bacterial or fungal abundance were normalized and expressed as percentage by dividing the speTSMs of each taxon by the total bacterial or fungal speTSMs within the sample. Proteins were grouped into protein families, and their abundance was evaluated by spectral counts after applying the parsimony principle.

### Functional analysis

Protein annotation with KEGG identifiers was carried out using the GhostKoala web service (https://www.kegg.jp/ghostkoala/) to match proteins with KEGG pathway maps. The spectrum count values for peptides associated with each KEGG pathway through protein mapping were summed to assign an abundance value to each functional term. Additionally, peptide-to-taxon mapping was performed to enable taxon-resolved functional quantification. The abundance of a function-genus association was normalized by weighting it with the ratio of the total functional signal of a given genus to its normalized abundance.

### Statistical analysis

Alpha-diversity (Shannon index) and beta-diversity (Bray-Curtis dissimilarity) were calculated using the vegan R package version 2.6-4 ^41^. Differential taxonomic and functional analyses were characterized using the fold change (FC). These results were statistically validated with the Welch t-test, which is used to compare the means of two groups with unequal variances. Differences between two conditions with an absolute fold change greater than 1.5 and a p-value less than 0.05 are considered statistically significant.

## Results

Overview of the experimental set-up for the proteotyping and metaproteomics analyses The influence of the diet on the gut microbiota of *G. fossarum* was established with animals subjected to the four different diets: Alder (reference), Carrot, Spinach, and Granules (rich in protein). The entire dataset acquired on pooled intestines of *G. fossarum* consisted of 1,708,964 MS/MS spectra, with 244,340 spectra reliably assigned to peptide sequences (14% attribution rate). The list of peptides detected per sample is provided in Table S1. The confident taxonomic proteotyping was based on 161,390 speTSMs reported at the genus taxonomical rank. As expected, they were attributed mainly to the host (152,364 speTSMs; 94.4%), the microbiota (6,288 speTSMs; 3.9%), and the residual food (2,738 speTSMs; 1.7%).

A total of 1,917 proteins were annotated, with 1,100 belonging to the host (57%), 607 to the microbiota (32%) and 210 to the residual food (11%). Table S2 summarizes the average number of speTSMs at the genus level and proteins assigned to host, microbiota, and residual food for each diet. The ratio of the microbial and residual food fractions depends on the diet. While the host/microbiota signals ratio was similar between the Alder diet and the Carrot diet (ratio = 14), that of the Spinach and Granules diets showed great differences (ratio = 28 and 71, respectively). Granules and Spinach diets were associated with lower microbial and residual food signals. To account for the varying microbial loads among animals with different diets, the data were normalized to express the abundance of microbial taxa as a percentage of the total microbial signals.

### The gut microbiota structure as a function of diet

The proteotyping analysis of the gut microbiota confirmed the presence of 37 bacterial genera and 5 fungal genera belonging to 5 and 2 phyla, respectively (Table S3). In all diet conditions, the gut microbiota was dominated by *Proteobacteria* (38%)*, Actinobacteria* (36%)*, Firmicutes* (12%) and *Bacteroidetes* (5%). *Cyanobacteria* were specific to the Carrot diet (Figure 1A). Fungi were represented by *Ascomycota* and *Oomycota,* with some variation in abundance across the different conditions (Figure 1A). The effect of the diet on the taxonomic composition was analyzed, with the Alder diet as the reference condition. While only few significant changes in abundance were observed at the phylum level, a strong diet effect was observed at the genus level (Figure 1B). To further assess the diversity of the microbial communities across different diets, alpha and beta diversity indexes were calculated based on the genus number and abundances. The Shannon index revealed a higher alpha diversity for the Spinach diet while no significant difference was observed for the other diets compared to the reference Alder diet (Figure S1A). The non-metric multidimensional scaling (nMDS) ordination of Bray-Curtis distances (Figure S1B) showed a clear separation of the different diets. These results confirm that microbial diversity tends to be impacted by the diet in terms of genus occurrence and abundance. Principal components analysis (PCA) performed at this taxonomical rank revealed clustering of the samples according to the diet (Figure 2A), indicative of a strong correlation between the short-term feeding diets and the microbial community structure. A low inter-individual variability was observed within the same diet. Figure 2B shows the specific and shared genera as a function of the diet. Between six to eight genera were unique to each diet, except for the Granules diet, which had none. Amongst the most abundant genera specific to a particular diet, *Actinoplanes* was accounting for 19% of the gut microbiota biomass in the animals fed with the Alder diet, *Flavobacterium* (17%) for the Spinach diet and the cyanobacterium *Gloeocapsa* (21%) for the Carrot diet. A total of 10 bacterial genera and 1 fungal genus were found to be common across all diet conditions. These 11 shared genera represent the most abundant microorganisms, constituting the core microbiota of *G. fossarum* and accounting for 49% of the total gut microbiota biomass. This core microbiota was dominated by *Streptomyces* (14% of the gut microbiota), following by *Clostridium* (6%), *Paenibacillus* (5%), *Sphingomonas* (5 %), *Pseudomonas* (4%), *Aspergillus* (3%), *Bacillus* (3%), *Acidovorax* (3%), *Flavobacterium* (2%), *Nocardia* (2%) and *Actinomadura* (1%). In addition to the presence or absence of certain genera, changes in the abundance of several genera also contribute to the differences observed between the different diets. Specific signatures of the most abundant genera for each diet were highlighted by a heatmap analysis (Figure S2). Compared to the reference Alder diet, 14 genera were significantly under-represented and 14 genera were over-represented in at least one diet (|FC| > 1.5; p-value < 0.05), while 14 genera were not significantly affected by the diet (Figure S2). The gut microbiota structure in *G. fossarum* showed the most pronounced changes when animals were fed with the Carrot and Granules diets, with 11 and 9 genera exhibiting lower abundances and, 8 and 5 genera with higher abundances, respectively. The microbial structure was also modified in the animals fed the Spinach diet, with the under-representation of 8 genera and the over-representation of 3 genera. Within the core microbiota, 7 genera were modulated in at least one diet condition, including 3 genera for the Carrot and Spinach diets, and 4 genera for the Granules diet. These less abundant modulations indicated greater stability in the structure of the core microbiota under different diets compared with the whole microbiota.

**Figure 1:**
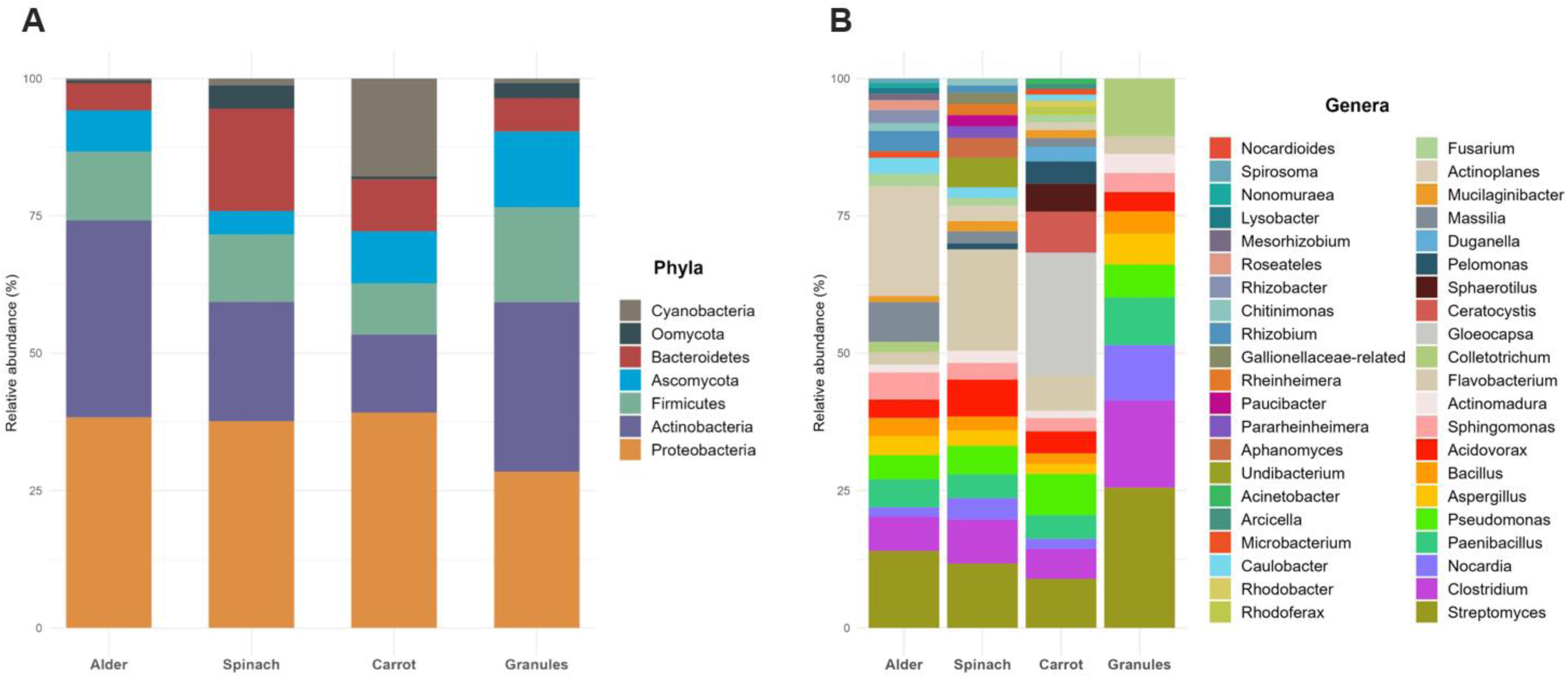
Gut microbiota composition of Gammarus fossarum depending on the diet expressed at (**A**) the phylum level or (**B**) the genus level.

**Figure 2:**
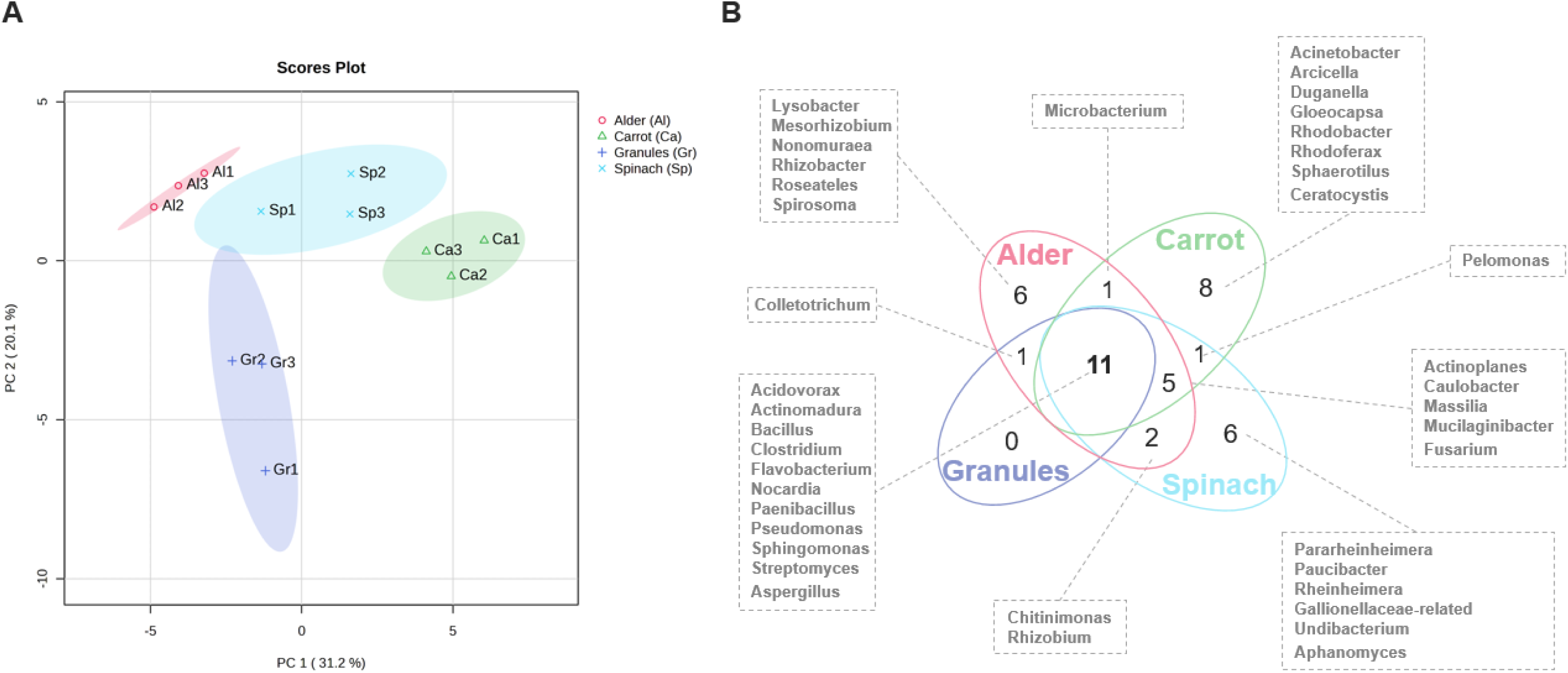
**(A)** Principal Component Analysis (PCA) score plot of the gut microbiota taxonomic composition, at the genus level, of the three replicates of Gammarus fossarum subjected to different diets. **(B)** Venn diagram showing the number of common and specific genera per diet identified and validated in the gut microbiota of Gammarus fossarum.

### Foodborne microbes contribute to the gut microbiota structure

In order to understand the contribution of microorganisms ingested by the animals, the food from the different diets was analyzed with the same metaproteomic pipeline. Proteotyping easily revealed the extent of colonization of food samples by bacteria and fungi. Spinach and alder leaves were the foods having the highest microorganisms/food ratio with 93 % and 78 % of microbial speTSM, respectively (Table S4). Although the number of microbial speTSM was high for the granules, 48 % of the signal was attributed to microorganisms (Table S4). The carrot had the lowest ratio with only 12 % of the speTSM related to microorganisms (Table S4). A reliable identification of 29 bacterial genera and 4 fungal genera was obtained for Alder leaves, while 35 bacterial genera and 4 fungal genera were identified for Spinach. Carrot exhibited 15 bacterial genera and 4 fungal genera, whereas Granules were shown to contain 18 bacterial genera and 6 fungal genera. Consequently, the high load of microorganisms within the food logically impacts the microbial structure of the gut after food ingestion. As highlighted in Figure 3, between 62 % and 92 % of genera identified in the gut microbiota for a given diet were also detected in the corresponding food. Conversely, microorganisms only detected in the gut could potentially be indigenous symbionts, such as *Microbacterium*, *Nonomuraea*, *Spirosoma*, *Mesorhizobium*, *Rhodobacter* and *Lysobacter*. Within the core microbiota, all microorganisms were detected in each of the analyzed foods, except for *Nocardia*, *Acidovorax* and *Sphingomonas* that colonized two or three foods only.

**Figure 3:**
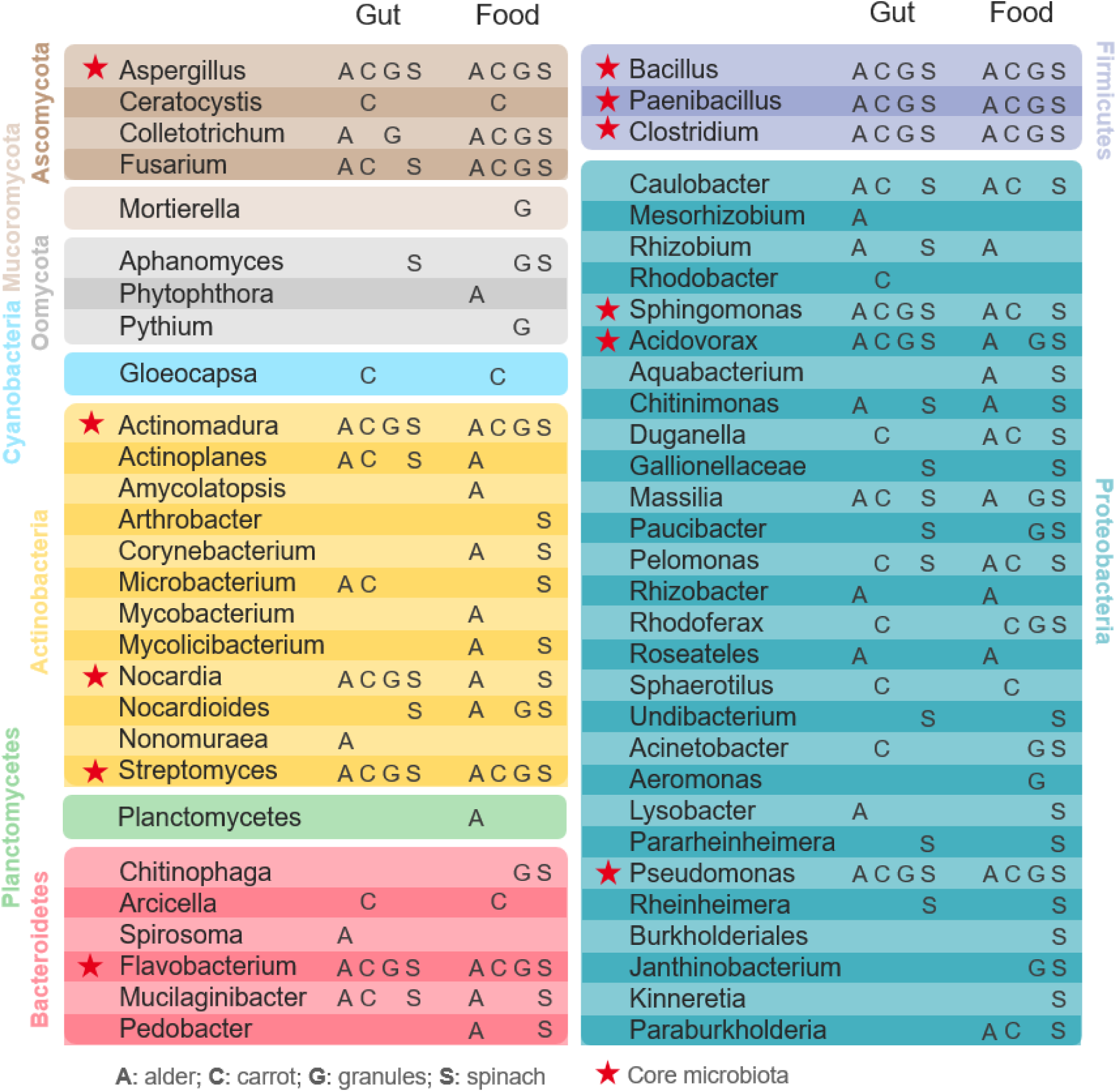
Comparison of microorganism occurrence in the gut microbiota versus the food.

### Gut microbiota function remains stable across diets

The gut microbiota function of *G. fossarum* fed with different diets was assessed by the mapping of annotated microbial proteins onto the KEGG Pathway database (Table S5). A total of 568 KEGG pathway-genus associations were listed. As expected, most microbial metaproteomic signals identified for the reference diet (Alder) are mainly involved in digestive processes. Carbohydrate metabolism, energy metabolism and signal transduction were accounting for 21%, 20%, and 12% of the functional signal quantities, respectively (Figure 4). Membrane transport (8%), transport and catabolism processes (7%), amino acid metabolism (5%) and metabolism of other amino acids (4%) were also significant (Figure 4). Fourteen additional pathways each represented less than 3% of the functional signal abundance (Figure 4). More detailed pathways are described in Table S6 with 1675 pathway-genus associations. For example, regarding the carbohydrate metabolism, the gut microbiota as a whole is mainly active throughout glycolysis, glyoxylate and dicarboxylate metabolism, starch and sucrose metabolism, galactose metabolism, pyruvate metabolism, and citrate cycle (Table S6).

**Figure 4:**
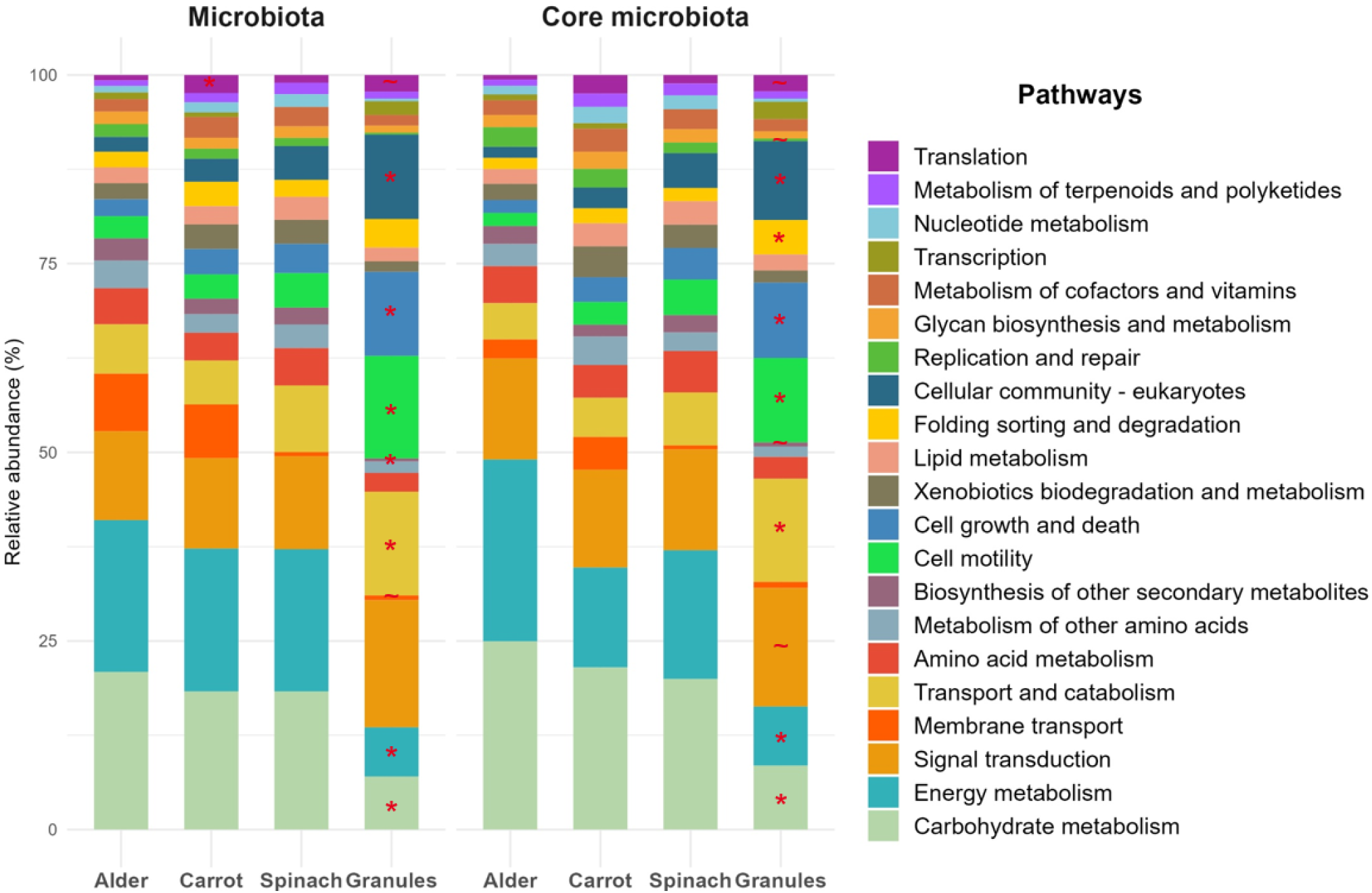
Main functions of proteins belonging to the gut microbiota or the core microbiota based on a KEGG pathway analysis. A significant modulation of a function compared to the reference Alder diet is highlighted by an asterisk * (|FC| > 2; p-value < 0.05) or a tilde ∼ (|FC| > 1.5; p-value < 0.05).

The investigation of functional changes across different feeding conditions, using Alder as the reference diet, revealed some striking differences. For example, only a few significant differences (|FC| > 1.5; p-value < 0.05) were observed for the Carrot diet, such as an increase in microbial translation (FC = 2). No change was observed for the Spinach diet (Figure 4). However, the Granules diet appeared to profoundly modify the functioning of the microbiota with nine functions significantly modulated (Figure 4). The main functions of the microbiota, such as carbohydrate metabolism and energy metabolism, were down-modulated (FC = -2.7 and -2.8, respectively) at the expense of an increase in transport and catabolism (FC = 2), and cellular processes (e.g. cell motility, cell growth and death, and cellular community – eukaryotes; FC > 3.7).

When considering the core microbiota only, the same functional pattern (Figure 4) was observed. The diets similarly impacted the functions of both the core microbiome and the global microbiome. For example, the Granules diet modified eleven functions compared to the Alder diet with a |FC| > 1.5, including the down-modulation of carbohydrate metabolism (FC = -1.9) and energy metabolism (FC = -1.9) as shown in Figure 4.

The genus-function associations indicated in Table S5 and Table S6 show which genera are responsible for a functional change in a given diet compared to the reference diet. As shown in Figure 5, *Aspergillus, Bacillus* and *Clostridium* were responsible of the main functional changes observed in the Granules diet in both the microbiota and core microbiota. Indeed, a significant down-modulation in the carbohydrate metabolism and energy metabolism of these three genera was highlighted (Figure 5). *Clostridium* was also associated with the up-modulation of transport and catabolism (FC = 2.7), along with *Colletotrichum* (FC = 2.5) and *Streptomyces* (FC = 2.1), as well as cellular processes (FC > 2.6). Although the glycan biosynthesis and metabolism was not significantly modulated at the whole microbiota level, its down-modulation was observed in the two fungi *Aspergillus* (FC = -3.3) and *Colletotrichum* (FC = - 2.7). It is also interesting to note that certain amino acid metabolic pathways were significantly up-modulated in the Granules diet, such as tryptophan metabolism (FC = 1.5), histidine metabolism (FC = 1.6), beta-alanine metabolism (FC = 1.6), lysine degradation (FC = 1.7) and valine, leucine and isoleucine degradation (FC = 1.7) as indicated in Table S6.

**Figure 5:**
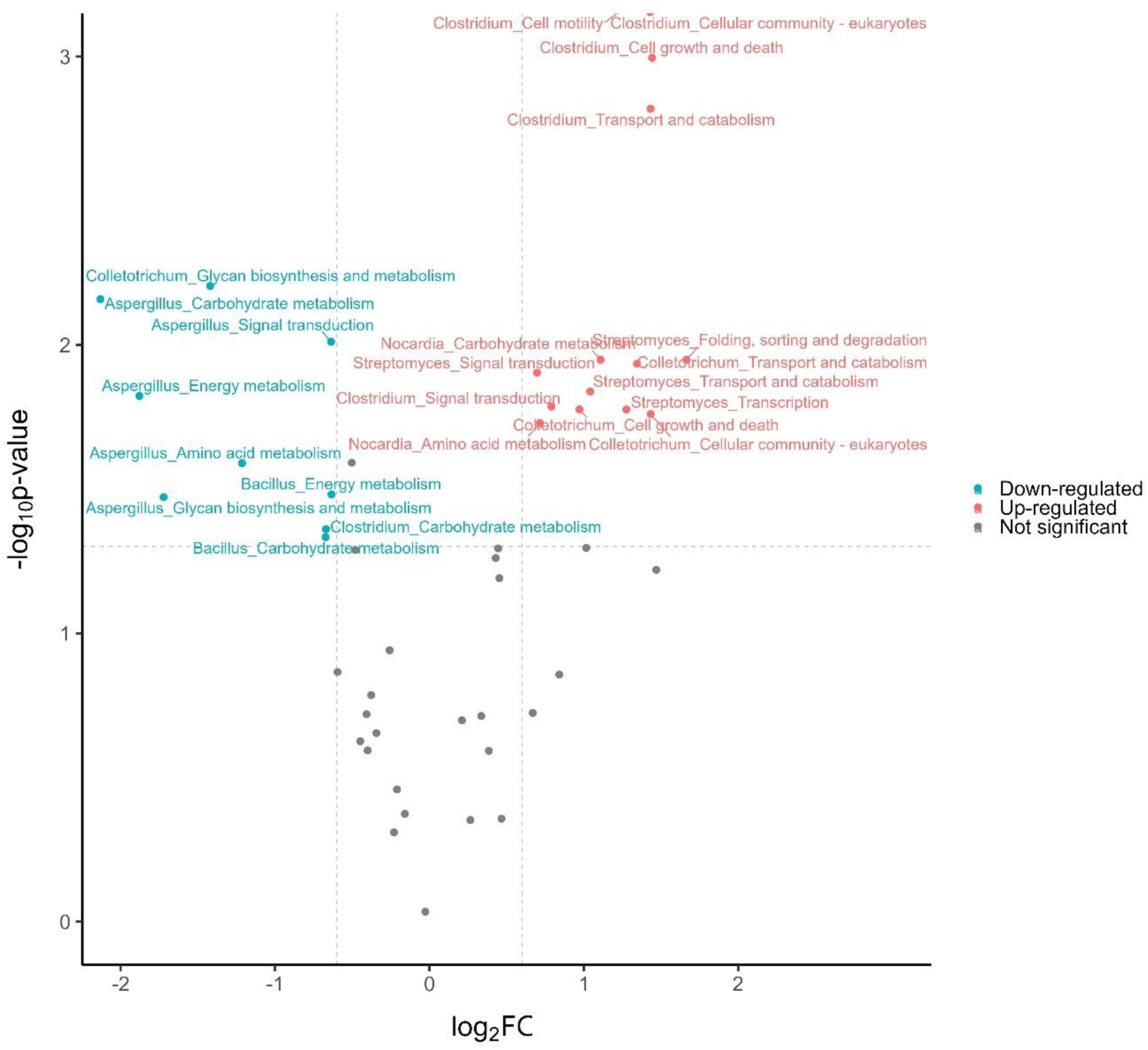
Volcano plot of the most abundant genus-function associations showing significant modulations (|FC| > 1.5; p-value < 0.05) for the Granules diet compared with the reference Alder diet.

To gain further understanding in the functional changes of the gut microbiota of *G. fossarum* fed with different diets, the most abundant microbial proteins were identified among the complete list given in Table S7. For each diet, the 10 most abundant proteins ranged between 1% and 6.4% of the recorded microbial protein biomass are presented in Figure 6. Several proteins were commonly found in every diet such as chitinase (mainly from *Paenibacillus*), beta-galactosidase (from *Paenibacillus*) and glyceraldehyde 3-phosphate dehydrogenase (from *Clostridium*, *Rhodoferax*, *Gloeocapsa*, *Pelomonas*, *Aspergillus* and *Fusarium*). In the microbiomes of the animals with Granules diet, seven of the most abundant proteins are of fungal origin (*Aspergillus* and *Colletotrichum*). These fungal proteins are involved in diverse processes such as in transcription and translation, carbohydrate metabolism, energy metabolism, signaling pathways, cell motility and transport (Figure 6).

**Figure 6:**
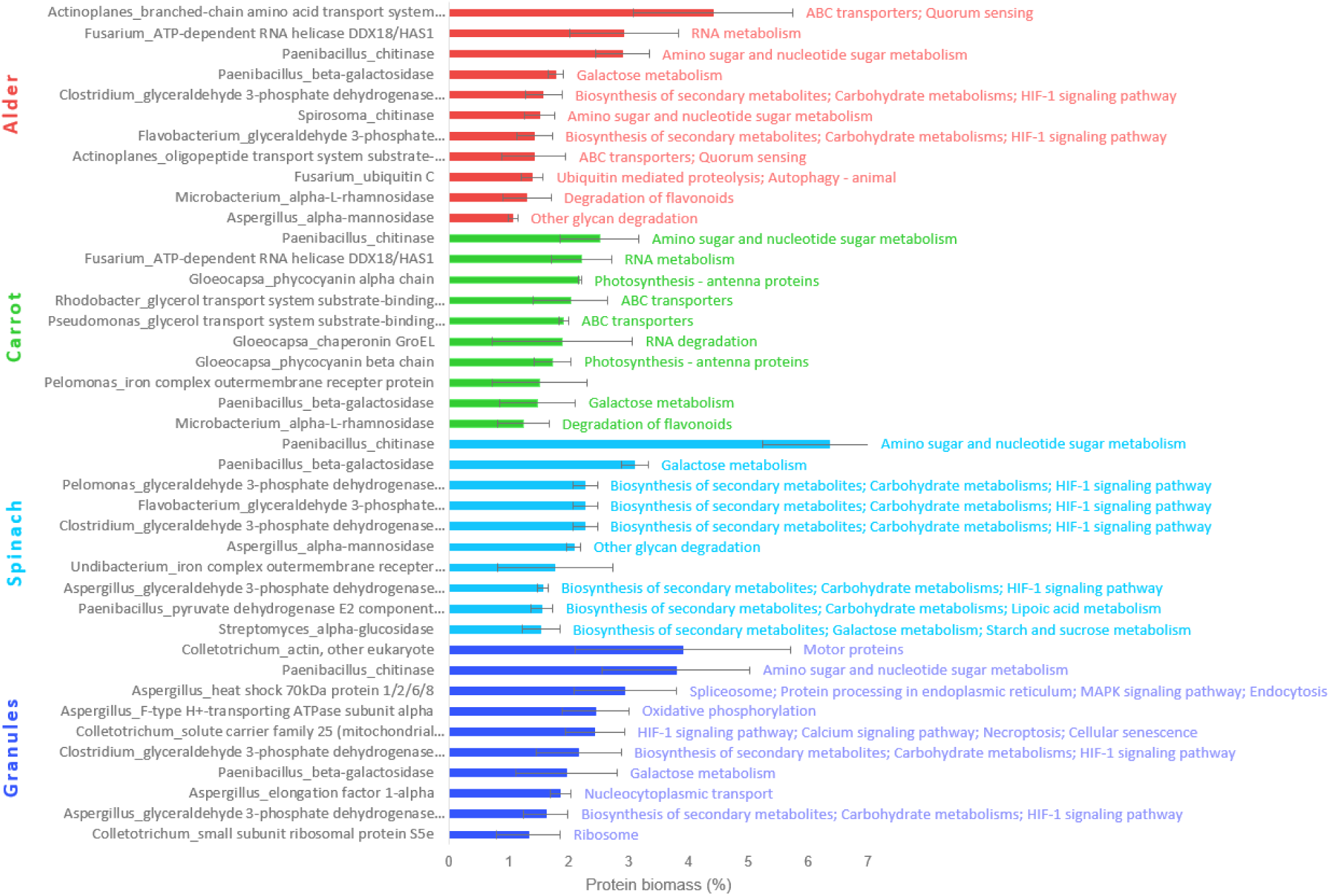
Most abundant microbial proteins, and their function(s), within the gut microbiota of Gammarus fossarum fed with different diets.

### Foodborne microbes contribute to the digestive effort of the gut microbiota

The viability of foodborne microorganisms in the animal gut was addressed by comparing their functional profile before and after ingestion (Table S8). Hereafter are two representative examples: the bacteria *Paenibacillus* and the fungi *Aspergillus* are both detected systematically in the different tested foods and the *G. fossarum* gut. The main metabolic functions of *Paenibacillus* were related to carbohydrate and energy metabolisms (e.g. carbon metabolism, glyoxylate and dicarboxylate metabolism, glycolysis, etc.) as well as to biosynthesis of secondary metabolites in both gut and food samples (Figure S3A). However, carbohydrate metabolism appeared to be more active in the gut with an increase of protein biomass involved in galactose metabolism, amino sugar and nucleotide sugar metabolism and starch and sucrose metabolism. Regarding the metabolic functions of *Aspergillus*, their profiles observed in the animal gut and the food samples were similar (Figure S3B). Biosynthesis of secondary metabolites and carbon metabolism were the two main metabolic pathways, followed by several pathways involved in carbohydrate and amino acid metabolism. *Aspergillus* also produced the enzyme cellulose 1,4-beta-cellobiosidase, as all the fungi found on alder leaves, allowing the degradation of cellulose (Table S9). However, once in the gut, this enzyme was no longer detected. According to their metabolic activity, foodborne microorganisms appeared to be viable in the intestine of *G. fossarum* during this short-term feeding diet, and contributed to the digestive effort of the gut microbiota.

### Diet effects on the host proteome

Notably, metaproteomics offers distinct advantages over other omics approaches in both identifying and quantifying host proteins in addition to the microbiome. The functional analysis of the identified host proteins provided insights into the metabolic activities and the molecular and cellular processes occurring in the intestine. (Table S10). Within the main host metabolic pathways for the reference Alder diet: carbohydrate metabolism represented the most abundant function in terms of host protein biomass (31.4%), followed by signal transduction (15.5%), transport and catabolism (9.9%), glycan biosynthesis and metabolism (8.7%), lipid metabolism (4.8%) and metabolism of other amino acids such as glutathione (4.7%) (Figure 7). A non-negligible protein biomass was also involved in host defense, especially immune system (3.1%), while the remaining 21.9% of the protein biomass was implicated in twenty other pathways, each accounting for less than 2.7% (Figure 7). Our results also highlight the most abundant protein from the enzymatic arsenal of *G. fossarum* (Table S11). Five of them represented more than 6% of the total protein biomass from the host, such as endoglucanase (16.2%), alpha-amylase (6.8%), chitinase (6.8%) and cellulose 1,4-beta-cellobiosidase (6.4%).

**Figure 7:**
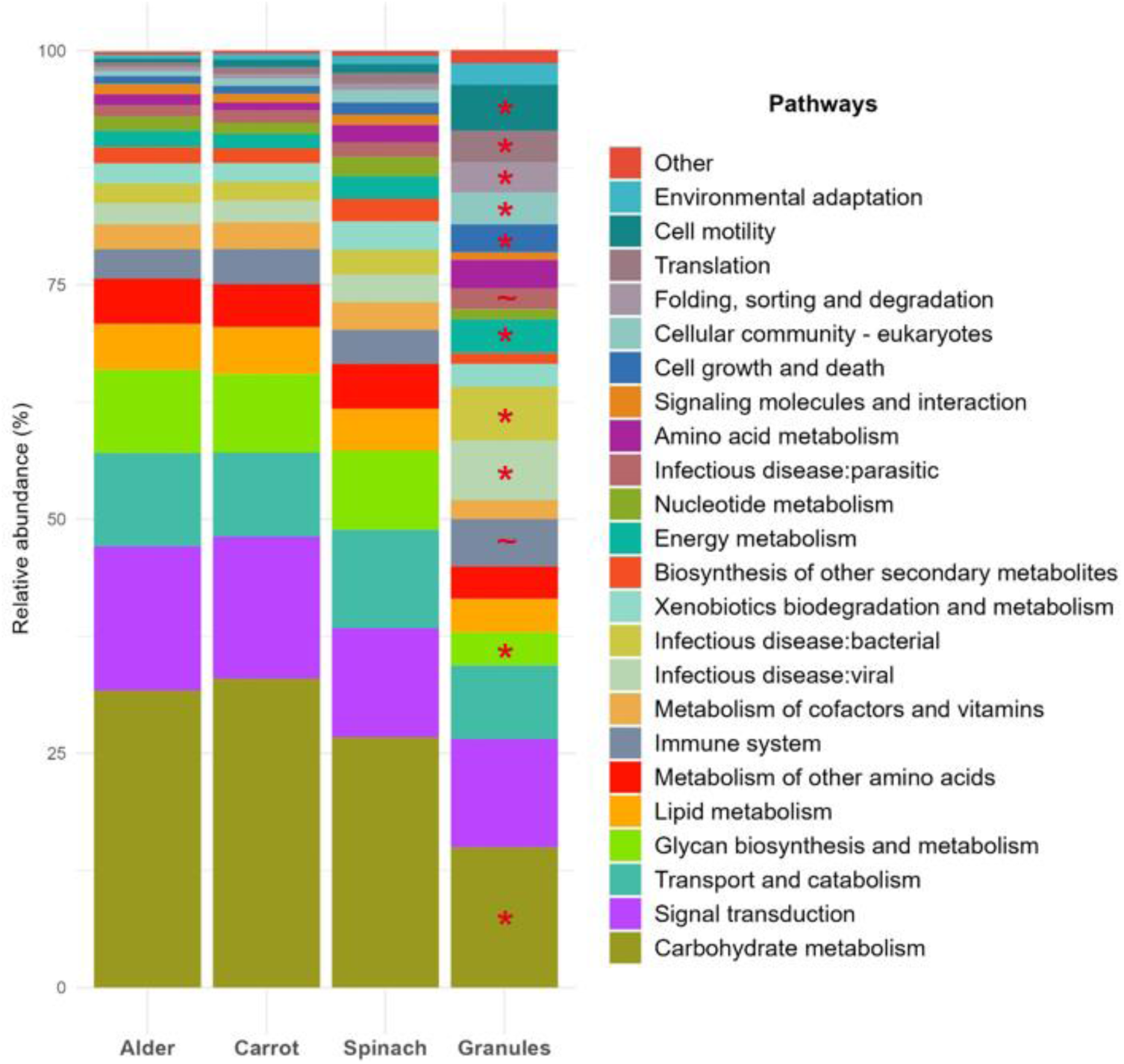
KEGG pathway analysis based on the abundance of host proteins identified for each diet. A significant modulation of a function compared to the reference Alder diet is highlighted by an asterisk * (|FC| > 2; p-value < 0.05) or a tilde ∼ (|FC| > 1.5; p-value < 0.05).

The study of the functional landscape of host proteins under different feeding conditions revealed significant changes in metabolic pathways and cellular processes for the Granules diet, while negligible or sparse modifications were observed for the Carrot and Spinach diets, respectively (Figure 7 and Table S10). Mainly, the carbohydrate metabolism (FC = -2.1) and glycan biosynthesis and metabolism (FC = -2.5) represented less important metabolic pathways for the host when it was fed with Granules, in full agreement with the high protein content of this food. Inversely, nine other pathways showed a significant increase in terms of protein biomass, including energy metabolism (FC = 2.1), cellular community (e.g. focal adhesion, adherens junctions, tight junctions and gap junctions; FC = 6) or cell motility (e.g. motor protein and regulation of actin cytoskeleton; FC = 11.3).

At the protein level, the main significant modulations were highlighted when comparing the most abundant host proteins found in the intestine of animals fed with Granules or Alder diets (Figure 8). Most of them were up-modulated in the Granules diet such as the enzymes carboxypeptidase Q (FC = 2.2), glutathione-S-transferase (FC = 6.5) or the structural proteins collagen type V/XI/XXIV/XXVII alpha (FC = 3), actin beta/gamma 1 (FC = 14.5) and tubulin beta (FC = 17). Finally, few proteins were down-modulated such as chlorophyllase (FC = -2.2) or alpha-N-acetylglucosaminidase (FC = -3.4), as well as the most abundant protein, endoglucanase, which tend to be down-modulated (FC = -2.3; p-value = 0.071).

**Figure 8:**
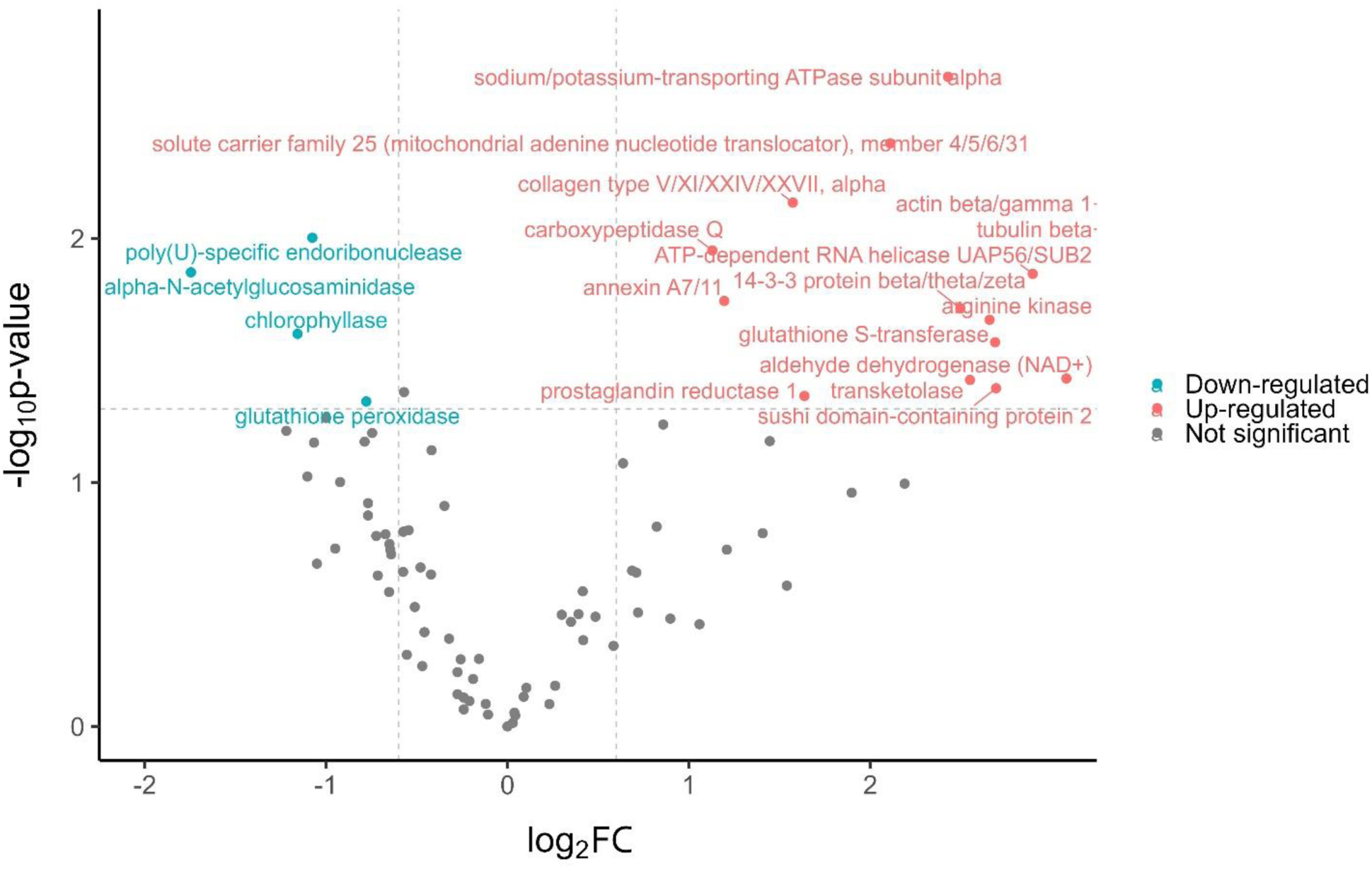
Volcano plot of the most abundant host proteins showing significant modulations (|FC| > 1.5; p-value < 0.05) for the Granules diet compared with the reference Alder diet.

Evidences also support an activation of the proteasome-ubiquitin system in the animals fed with the Granules diet. These comprise the detection of 33 proteins related to the 20S and 26S proteasome that were absent in the reference diet, showing a unique role for this complex in recycling foodborne proteins in the intestine (Table S11). Although ubiquitin C was down-modulated (FC = -8.7), for the Granules we detected ubiquitin-small subunit ribosomal protein S27Ae, ubiquitin-conjugating enzyme E2 N, small ubiquitin-related modifier and ubiquitin-activating enzyme E1 that were absent in the reference diet (Table S11).

## Discussion

In this study, metaproteomics gains unique insights into the digestive system of *G. fossarum* by assessing the changes of the gut microbial communities, their functions, and the host proteome, depending on four short-term feeding conditions. The data gleaned from the metaproteomic analysis of the intestinal microbiome broadened our understanding of host-microbiota interactions within the digestive system of gammarids and by extension, amphipods in general. The gut microbiota of *G. fossarum* is primarily composed of four dominant phyla, listed in descending order of abundance: Proteobacteria, Actinobacteria, Firmicutes and Bacteroidetes. Here, we identified the 42 most abundant microbial genera (37 bacteria and 5 fungi) in the gut microbiota of *G. fossarum* subjected to different diets. In line with the literature, 37 of these genera have already been identified in the gut microbiota of amphipods ^31,42–47^ and/or in beech or alder leaf litter, representing the main food resource of these detritivorous animals ^48,49^. The consistent alignment of our metaproteomic findings with existing literature lends weight to our methodology given that most current studies primarily rely on nucleic acid sequencing techniques, such as 16S rRNA amplicon sequencing and metagenomics. To our knowledge, we also reported for the first time the presence of the Actinobacteria *Nonomuraea* and the Proteobacteria *Rhizobacter* in the gut microbiota of an amphipod, highlighting possible specific genera depending on the species under scrutiny. However, other organisms have been reported to harbor these bacteria in their intestines, such as zebrafish and bees ^50–52^. Hence, the dominant microbial genera hosted in the intestines of *G. fossarum* are commonly found in the gut microbiota of other animals, including amphipods. We anticipate that a broader array of host-specific microorganisms could be observed if a finer discrimination at the species level is obtained. For this, more extensive datasets should be recorded by in-depth nucleic acid sequencing approach or next-generation tandem mass spectrometry methodology. As shown recently, reaching this taxonomical rank is feasible with the latest Orbitrap Astral instrumentation, which enables the recording of high quality MS/MS data at remarkable speed ^11^. However, achieving a more comprehensive analysis would necessitate a greater quantity of biological material.

The microbiota influences various host functions, including digestion, development, metabolism, immune defenses, and behavior, forming an integrated functional entity known as the “metaorganism” ^53,54^ or “holobiont” ^55,56^. In invertebrates, an increasing body of evidence is unraveling the mechanisms that govern the intricate host-microbiota interactions ^57,58^. In this study, invaluable information on the functionality of the gut microbiota in *G. fossarum* under normal conditions (Alder diet) were obtained by directly assessing the proteins, the workhorses of biological system metabolism, and their abundances, which serve as proxies for their activity levels. As expected, both the host and the microbiota possess a large arsenal of enzymes for breaking down decomposing organic matter. They both actively participate in digestive processes and nutrition, as shown by carbohydrate metabolism and, to a greater extent for the microbiota, energy metabolism (e.g. oxidative phosphorylation). Our results are in line with the functions of the gut microbiota depicted in three hadal amphipods, a group of crustaceans inhabiting the deepest parts of the aquatic compartment, thus adapted to survive extreme conditions ^43^. Their study based only on the gene catalogue established by metagenomics has shown that genes encoding proteins involved in carbohydrate metabolism are indicating the most enriched metabolic pathway, including glycolysis, amino sugar and nucleotide sugar metabolism, starch and sucrose metabolism, and pyruvate metabolism ^43^. The food consumed by *G. fossarum,* such as decomposing alder leaves, contain high levels of various carbohydrates such as free sugars or more complex polysaccharides (i.e. glycans) like cellulose, pectin or hemicellulose. While both the microbiota and the host metabolized free sugars, we observed that the degradation of certain glycans, such as cellulose, is mainly performed by the host. Indeed, this stands as a significant advantage of metaproteomics over other omics methodologies, as it allows for the analysis of the host’s enzymatic repertoire. For instance, endoglucanase and cellulose 1,4-beta-cellobiosidase, enzymes involved in cellulose degradation, rank among the top 5 most abundant proteins from the host. However, endoglucanase has been detected only in trace amounts in *Bacillus* and *Paenibacillus*, while cellulose 1,4-beta-cellobiosidase was exclusively found in fungi colonizing alder leaves. Three decades ago, numerous studies have explored the cellulose degradation potential of *Gammarus* species through microbial or endogenous host enzymes. However, the experimental methodologies of that time were constrained, leading to conflicting conclusions ^59,60^. Nevertheless, most evidence supports our view that the predominant role in cellulose degradation is played by endogenous enzyme activity, with microbial enzymes playing a secondary contribution ^61^. Over the past two decades, growing evidence has shown that some animals, including aquatic invertebrates, possess genes that code for endogenous cellulases. ^62^. Our metaproteomic study contributes to these discoveries, particularly in the understudied group of amphipods, by proving the synthesis of these host cellulases. Otherwise, other glycans, such as chitin, are cooperatively degraded by both the host and the microbiota. We observed a high level of chitinase originating not only from the host but also from two bacteria, *Paenibacillus*, already recognized for its significant production ^63^, and *Spirosoma*. Chitin, among the most abundant organic compounds in nature along with cellulose, is a structural component found notably in the cell walls of fungi and the exoskeletons (e.g. cuticle) of arthropods ^64^. It is well known that *Gammarus* spp. show preferences for substrates that has been previously colonized by fungi, thereby increasing the nutritional value of the food ^32^. We provide evidence that the host-microbiota system is well-adapted to the digestion of foodborne fungi by chitinase, as a valuable nutrient source. Hence, fungal chitin can be degraded, as well as the chitin from the old cuticle of the host can be recycled during the molting process of the host ^64^. Therefore, our results demonstrate the essential role of the gut microbiota of *G. fossarum* and its functional complementarity with the host. Furthermore, our findings highlight the importance of signal transduction, raising the question of possible communication between the microbiota and the host. This process ranks as the third most abundant for the microbiota and the second most abundant for the host. Signal transduction plays a crucial role in regulating the immune system, metabolism, hormones, neurotransmitters, as well as tissue development and repair ^58,65^. The biosynthesis of secondary metabolites by gut microbes, as represented in this study through related proteins, can interfere with host signaling pathways, affecting host physiology and developmental processes in various ways, as previously mentioned ^65^.

In our study, the impact of short-term diets on the microbial community resulted in a significant reorganization of taxonomic composition in some cases, particularly notable in the transition from an Alder leaves diet to a protein-based diet (Granules). However, a core microbiota of eleven genera was maintained as a common base. Two possible explanations can justify the changes in microbial structure. First, the food is colonized by microorganisms, which are ingested with the food and remain at least temporarily viable in the animal gut. Hence, food-associated bacteria and fungi can modify the profile of gut microbiota. Secondly, the observed changes in the microbial community after feeding may also stem from less abundant bacterial strains that, despite being present in the gut, go undetected unless their prevalence is reinforced by particularly beneficial feeding and growth conditions.

Based on our results, the ingestion of foodborne microorganisms contributes to short-term structural changes in the gut microbiota of gammarids. Until now, the scientific literature addressing the connection between foodborne and gut microbiome focuses mainly on human subjects. Although the mechanisms governing these interactions are complex and sparsely understood, studies already demonstrated that foodborne microbes can transiently colonize the human gut and rapidly cause major shifts in the microbial composition ^66–68^. A recent metagenomics study revealed that foodborne microbes represent on average up to 3% of the gut microbiome of adult human ^69^. To our knowledge, the shaping of the gut microbiota by foodborne microorganisms has not yet been directly demonstrated in gammarids, amphipods or shrimps, but their detritivorous lifestyle argues for a strong impact of dietary microbiota. It is well known that microorganisms present in the surrounding environment, such as water or sediment, can transit in the gut of aquatic organisms. For instance, several studies revealed a high similarity of bacterial communities between the gut of shrimps and the surrounding sediment and water ^70–73^. In this study, the water used for culturing gammarids and for decomposing food might contain ubiquitous microorganisms that were detected in both the food samples and the gut samples. This raises the question of whether the core microbiota originates from the food, the water, or was already present in the gut. For gammarids, long-term studies could be conceived to highlight whether the colonizing foodborne microorganisms will persist in the gut or are merely transient, but this would need an improved precision at the species, subspecies, or even strain level that can be obtained only with more in-depth research. It is essential to emphasize that the gut microbiota of detritivorous aquatic animals is probably more flexible than that of terrestrial organisms and is highly sensitive to environmental factors such as changes in diet as previously reported ^74^. Noteworthy, we did not differentiate which of the microorganisms are autochthonous or indigenous, *i.e.* able to colonize the gammarid’s intestinal epithelial surface, or allochthonous or transient, *i.e.* present only in the lumen. Additional experiments based on fluorescence in situ hybridization of probes specific of the identified microorganisms and microscopy could be planned in future projects to better delineate their spatial localization in the intestine and role.

The nutrients in food can create favorable gut conditions for the growth of certain microorganisms or the modification of their metabolism as well as that of the host. As highlighted in this study, most functional changes were observed for the microbiome of animals with the Granules diet, which is importantly enriched in animal and vegetal proteins compared to the other diets. This specific diet exerts measurable effects on microbial metabolism by activating certain amino acid metabolic pathways (e.g. tryptophan, histidine, lysine, beta-alanine, etc.) closely associated with protein degradation. A similar pattern has been observed in the gut microbiota of humans fed with a protein-rich diet ^75^. On the host side, the protein-rich granule content increased the abundance of certain peptidases (e.g. carboxypeptidase Q and Xaa-Pro dipeptidase), that play a role in peptide hydrolysis and protein degradation ^76,77^. This amino acid nutrient source can be used as cellular fuel, evidenced by an increase in energy metabolism in the host. In shrimp farming, the protein content in the diet plays a crucial role for promoting rapid growth of animals ^78^. Here, the granules provided to gammarids, originally designed for shrimp farming, are rich in proteins (e.g. α and β hemoglobin subunits, serum albumin, and myosin from *Sus crofa*) and increase protein catabolism by both the host and its microbiota. In the present work, functional analysis of host proteins provides evidence that these granules may contribute to the stimulation of gammarid growth. This is underpinned by the up-regulation of numerous processes integral to the structuration of cells and tissues. Notably, various aspects of cellular community dynamics, including focal adhesion, adherens junctions, tight junctions, and gap junctions, undergo up-regulation. In addition, cell motility (e.g. motor protein and regulation of actin cytoskeleton) and cellular growth and death processes were also up-modulated. Finally, the activation of proteasome observed in this study and the up-regulation of ubiquitin related proteins indicate the necessity of preserving cellular homeostasis through the regulation of protein turnover.

The proteasome-ubiquitin system is involved in the regulation of many basic cellular processes such as cell cycle and division, differentiation and development, the response to stress, DNA repair, regulation of the immune and inflammatory responses, apoptosis, and many others ^79,80^. Therefore, the inclusion of protein-rich granules in the diet not only enhances protein digestion with the assistance of microbiota but also initiates a cascade of molecular and cellular events that may promote the growth of *G. fossarum.* A proteomics study conducted on the Y-organ of the crab *Gecarcinus lateralis* substantiates our hypothesis by revealing the significance of cytoskeletal proteins (e.g. actin and tubulin) and ubiquitin-proteasome system in the molting process of this crustacean ^81^. The very different nutrients provided by granules have other consequences on the metabolism of the host and the gut microbiota. The carbohydrate metabolism that predominates in vegetable- or leaf-based diets loses its significance in the Granules diet, observed for both the host and its microbiota. Among complex carbohydrates, more abundant in plants, the glycan biosynthesis and metabolism was significantly down-modulated in the host and the two fungi *Aspergillus* and *Colletotrichum*. For instance, a down-regulation of alpha-N-acetylglucosaminidase, involved in the degradation of chitin, or a trend toward down-regulation of the most abundant protein, endoglucanase, which degrades cellulose, have been observed in the host. In line with the composition of the granules, we also observed a down-regulation of chlorophyllase, which catalyses the first step of the chlorophyll degradation, commonly present in plants and absent from the granules. Consequently, the main effects induced by the granules during a short-term diet lie in adapting the metabolism of the host and its microbiota to nutrients that are very different from those found in alder leaves or in the two other vegetable-based diets.

Our study delineated a core gut microbiota of 11 genera, representing 49% of the microbial protein biomass. Interestingly, the different diets had relatively little impact on the structure of this core microbiota in terms of genus abundance. In addition, the core microbiota exhibits analogous primary functions to those of the gut microbiota and, to a comparable degree, manifests a similar functional landscape across varying diets. The core microbiome is known to play a crucial role in sustaining host health, exerting a significant influence on the microbial community ^82^. In recent years, a growing interest in the core microbiome of cultured crustaceans has emerged with the aim of enhancing our understanding of host-microbiota interactions and, consequently, improving animal health ^16,72^. In their study, Jiang et al. (2023) observed a positive correlation between the relative abundance of core genera, dominated by *Photobacterium*, and the stability of the gut microbial community in the mud crab *Scylla paramamosain*. *Photobacterium*, which has probiotic activity, may play a critical role in enhancing the immune response of the mud crab and preserving the diversity of its gut microbiota. In our study, *Streptomyces* exhibits the highest abundance of the core microbiota and is known to have probiotic activity. For instance, Mazón-Suástegui et al. (2020) demonstrated the modulating effect of a *Streptomyces* strain on *L. vannamei* microbiota, along with its stimulatory effect on *Bacteriovorax* population and various antimicrobial producers that protected shrimp from *Vibrio parahaemolyticus* infection. Interestingly, biosynthesis of secondary metabolites is the most abundant functional pathway of *Streptomyces* associated to *Gammarus* gut microbiota, and one of the most abundant among all the genera-pathway (Table S4). Secondary metabolites can interact with the microbial community and the host, and are likely to stimulate growth and immune system. Therefore, the core genera and particularly *Streptomyces* could have beneficial impacts on the gut microbiota and the health of *G. fossarum*.

Studies investigating the impact of feeding on aquatic crustaceans have mainly focused on shrimps raised for human consumption ^13^. These experiments, spanning several weeks, have consistently demonstrated the diet’s influence on the gut microbiota composition, with potential repercussions for the host in terms of growth, digestive enzyme activity, immune system and pathogen resistance. While the primary focus of research has been on examining the impact of the diet on microbial structure, only a few studies have incorporated a more holistic approach to predict the functional aspects of the gut microbiota ^85,86^. However, it is essential to note that these studies, designed to enhance shrimp production, may not entirely mirror the natural conditions. Comparatively, microbiota from other non-commercial aquatic crustaceans, aside from some research on the model organism *Daphnia magna*, remains relatively less characterized. Host-microbiota interactions in *D. magna* have been found to be highly dependent on food availability ^87^. Another study on this Branchiopod crustacean indicated positive correlations between diet quality and susceptibility to antibiotic exposure ^88^. The consensus in the majority of studies suggests a considerable susceptibility of gut microbiota to changes in the host’s diet over the long term. In line with this trend, our findings reveal that the structure of the gut microbiota in *G. fossarum* can be influenced by various short-term diets. Importantly we have indicated for the first time that foodborne microorganisms have a significant impact on the microbial community of gammarids. Despite the reshaping of the gut microbiota, its functions remain stable, as observed in the case of the Carrot and Spinach diets. However, a particularly different diet, rich in protein, prompted a metabolic adaptation of the host and its microbiota to a new nutrient source. Therefore, differences in food preferences among species may influence the conditioning of gut microbial structures through foodborne microorganisms. This hypothesis could explain the species-specific bacterial communities observed in the guts of different talitrid amphipods ^42^.

Beside the novel insights gained into host-microbiota system of amphipods subjected to various diets, our study is also important in terms of ecotoxicology as gammarids are used as sentinel of the aquatic environment ^28,89^. As recently demonstrated, the gut microbiota plays a role in the response of organisms to contaminants ^90,91^. Our results suggest that the nutrients present in the exposure media should not significantly influence the function of the gut microbiota, as drastically different diets from one site to another, such as a high-protein diet, should be less frequent in the river litter. Otherwise, the modulations observed in this study could be used as a reference to discriminate the effects of diet variation from the variations measured in individuals collected at multiple sites in an ecotoxicological context. Particular attention should also be paid to the core microbiota because, given its importance, its disruption by contaminants could have consequences for the health of both the gut microbiota and the host. Ultimately, our metaproteomics approach opens up new possibilities for enhancing our understanding of host-microbiota interactions in response to environmental or intrinsic factors in non-model organisms, despite the challenges posed by the limited biomass obtained from millimetric animals and the complexity arising from the presence of intestinal tissues mixed with the microbiota.

In conclusion, the metaproteomics approach applied here brings experimental functional information on the gut microbiota of *G. fossarum* together with an interesting pan-kingdom taxonomical characterization, a unique feature offered by protein sequencing as recently discussed ^5^. Moreover, microbiome taxonomical and functional changes under different short-term feeding diets can be easily monitored. We have shown that the host and its microbiota have complementary metabolic activities allowing the degradation of complex polysaccharides, such as cellulose and chitin, but also exhibit a high level of signal transduction, that should deserve more attention in the future to improve our understanding of the holobiont interactions. While diet is determinant in the shaping of the microbial community structure, we demonstrate for the first time in amphipods that foodborne microorganisms strongly influence structural changes during short-term feeding. Evidence also support that foodborne microorganisms in transit in the gut were still viable after ingestion and contribute to the digestive effort of the gut microbiota. Functionalities of the host-microbiota system remain globally stable under the different diets. However, the protein-based diet induced functional shift in both the host and its microbiota due to their adaptation to a completely new source of nutrients. Our study also reveals the presence of a core microbiota driving the main functions of the gut microbiota, which appears to be less susceptible to diet variation. These findings hold significance for forthcoming ecotoxicological and biomonitoring investigations, which could leverage the microbiomes of these sentinel animals as pivotal targets.

## Data availability

The mass spectrometry proteomics data have been deposited to the ProteomeXchange Consortium via the PRIDE partner repository with the dataset identifier PXD055541 and 10.6019/PXD055541.

## Acknowledgement

This research was funded by the French *Agence Nationale de la Recherche* (Dyn-Microbiome project, ANR-20-CE34-0012) and the Région Occitanie Pyrénées-Méditerranée (DeepMicro grant). This work benefited from the French GDR “Aquatic Ecotoxicology” framework which aims at fostering stimulating scientific discussions and collaborations for more integrative approaches. JA expresses gratitude to the Région Occitanie, the French IBISA GIS network, the INBS ProFi, and the Agence Nationale de la Recherche for their invaluable support in advancing metaproteomics within the ProGénoMix platform.

## Author contributions

**TD**: Formal analysis, Investigation, Writing - Original Draft; **OP**: Methodology, Data curation, Writing - Review & Editing; **LG**: Methodology, Writing - Review & Editing; **DDE**: Writing – review & editing; **ND**: Resources; **OG**: Writing – review & editing; **AC**: Conceptualization, Writing - Review & Editing; **JA**: Conceptualization, Writing - Review & Editing, Funding acquisition

