## Supplementary figures and images for "Key gut microbiota components and functions in an aquatic keystone species across diets assessed by metaproteomics"

### Supplementary data (Figures S1-S3)

#### Slide 1
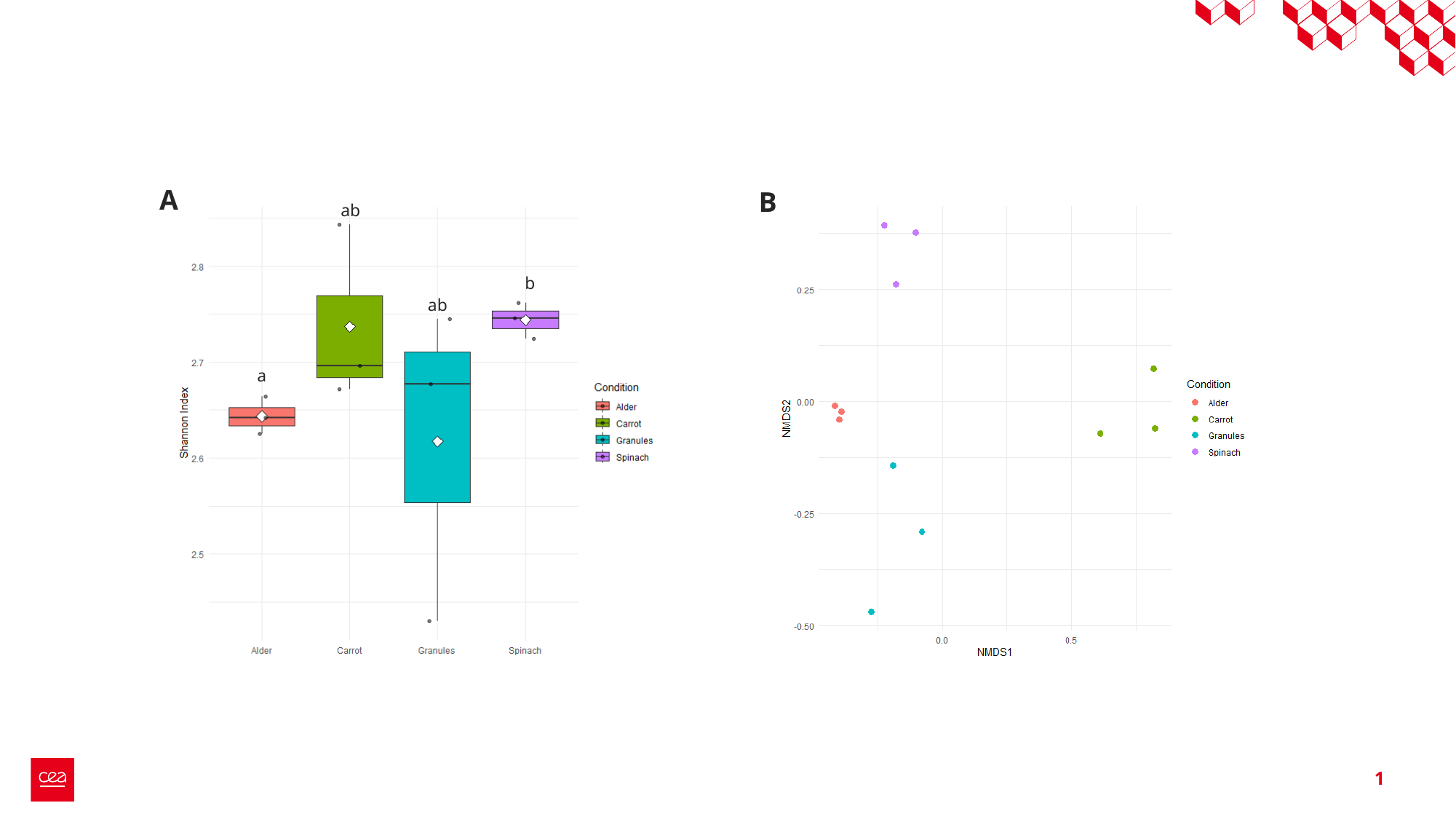

A
B
ab
b
ab
a
1

#### Slide 2
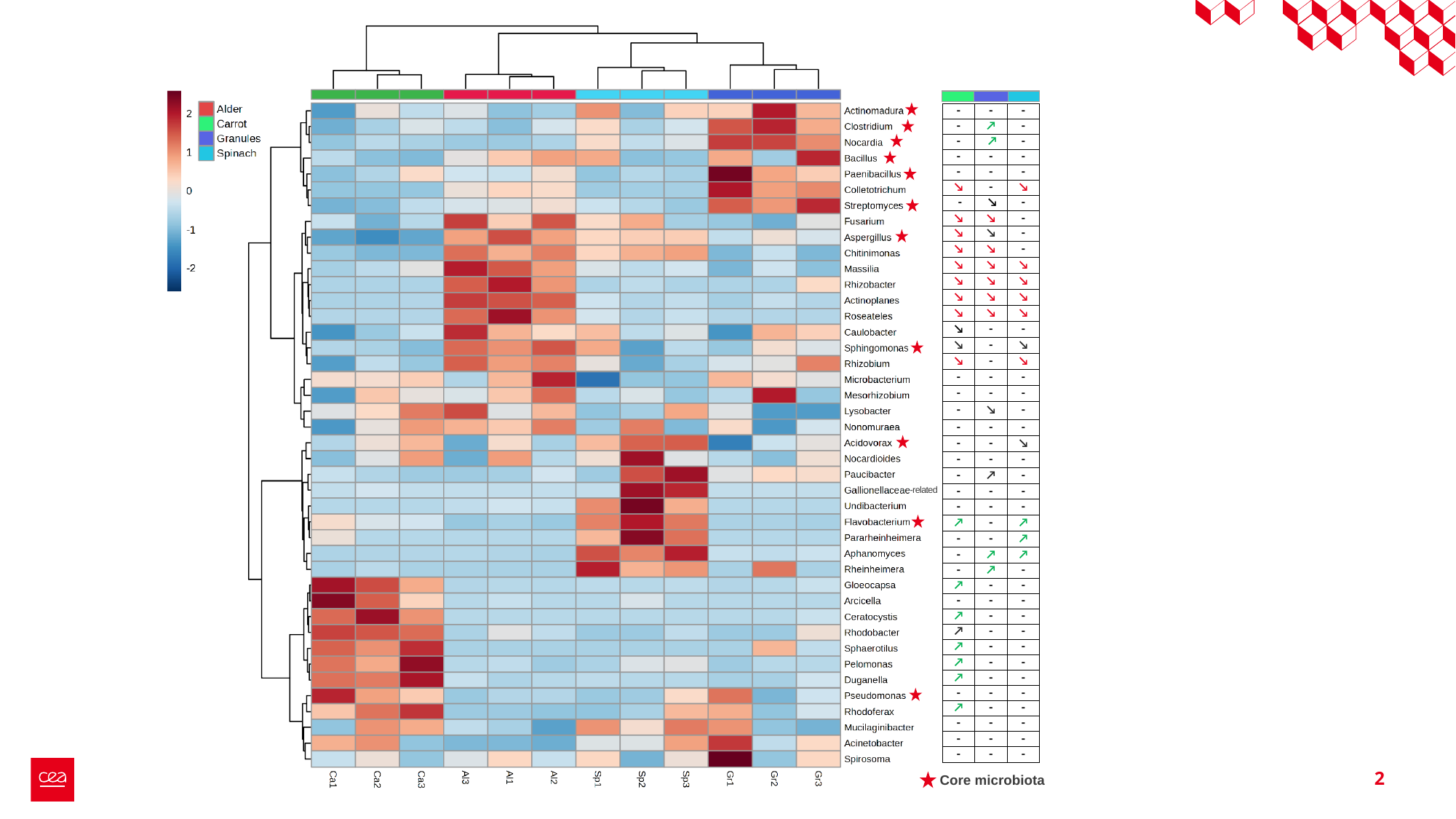

| - | - | - |
| --- | --- | --- |
| - | ↗ | - |
| - | ↗ | - |
| - | - | - |
| - | - | - |
| ↘ | - | ↘ |
| - | ↘ | - |
| ↘ | ↘ | - |
| ↘ | ↘ | - |
| ↘ | ↘ | - |
| ↘ | ↘ | ↘ |
| ↘ | ↘ | ↘ |
| ↘ | ↘ | ↘ |
| ↘ | ↘ | ↘ |
| ↘ | - | - |
| ↘ | - | ↘ |
| ↘ | - | ↘ |
| - | - | - |
| - | - | - |
| - | ↘ | - |
| - | - | - |
| - | - | ↘ |
| - | - | - |
| - | ↗ | - |
| - | - | - |
| - | - | - |
| ↗ | - | ↗ |
| - | - | ↗ |
| - | ↗ | ↗ |
| - | ↗ | - |
| ↗ | - | - |
| - | - | - |
| ↗ | - | - |
| ↗ | - | - |
| ↗ | - | - |
| ↗ | - | - |
| ↗ | - | - |
| - | - | - |
| ↗ | - | - |
| - | - | - |
| - | - | - |
| - | - | - |
-related
2
Core microbiota

#### Slide 3
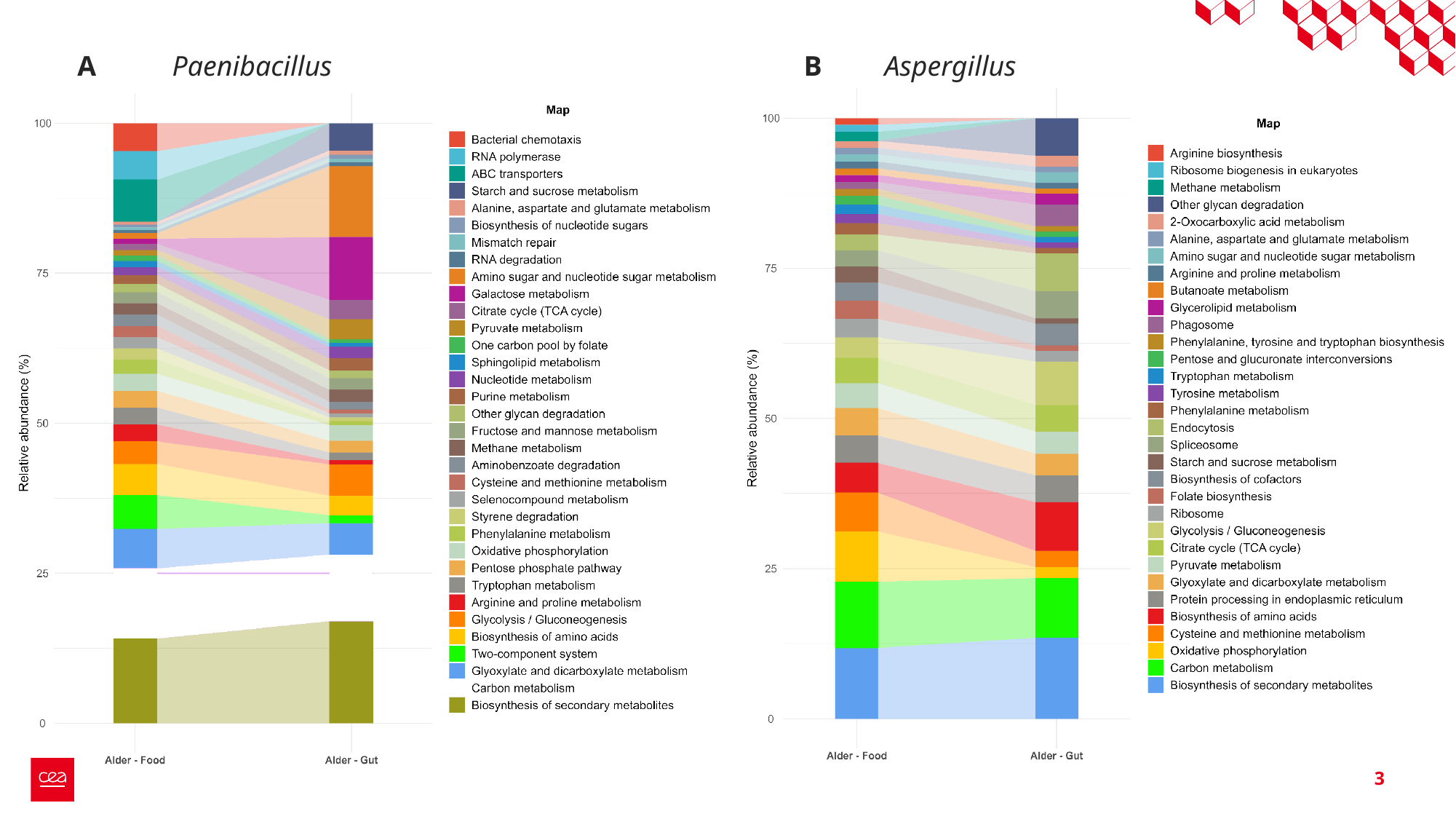

A
Paenibacillus
B
Aspergillus
3
